# Mpf1 is a novel factor that affects the dual distribution of tail-anchored proteins between mitochondria and peroxisomes

**DOI:** 10.1101/2024.04.30.591829

**Authors:** Nitya Aravindan, Daniela G. Vitali, Jessica Oberst, Einat Zalckvar, Maya Schuldiner, Doron Rapaport

## Abstract

Over half of cellular proteins must target upon their translation to distinct cellular compartments to function. Whereas considerable progress has been made in our understanding of targeting to individual organelles, we know truly little about how proteins distribute their targeting between two, or more, destinations – a process called dual/multiple targeting. In this study, we shine mechanistic insight into the process of dual targeting of proteins between the yeast mitochondria and peroxisomes. We performed a high throughput systematic microscopy screen in which we visualized the location of the model tail-anchored (TA) proteins Fis1 and Gem1 on the background of mutants in all yeast genes. This screen identified three proteins, whose absence resulted in a higher portion of the TA proteins in peroxisomes: the two paralogues Tom70, Tom71, as well as the uncharacterized gene *YNL144C* that we renamed <u>m</u>itochondria <u>p</u>eroxisomes factor 1 (Mpf1). We characterized Mpf1 to be an unstable protein that associated with the cytosolic face of the mitochondrial outer membrane. Furthermore, our study uncovers a unique contribution of Tom71 for the regulation of the dual targeting and reveals a link between *TOM70/71* and the quantity of transcripts of *MPF1*. Collectively, our study revealed, for the first-time, factors that influence the dual targeting of proteins between mitochondria and peroxisomes.

## Introduction

Eukaryotic cells have evolved complex machineries that direct, with the help of targeting signals, cytosolically synthesized proteins to specific intracellular locations. While most proteins target to a single compartment, there are those that are targeted to two (or even more) cellular destinations. Some examples of such proteins include the metabolic enzymes fumarase and aconitase, where in addition to the mitochondrial population a certain portion of the protein is also found in the cytosol or even the nucleus (Regev-Rudzki & Pines, 2007; Stein et al., 1994; Yogev et al., 2011).

Several cases of such dually targeted membrane proteins are those that distribute between mitochondria and peroxisomes. These two organelles maintain extensive crosstalk and are transiently associated by multiple contact sites. Pex11, a key protein involved in initiating peroxisome division was found to interact with Mdm34, a constituent of the ER-mitochondria encounter structure (ERMES) complex, thus mediating these mitochondria-peroxisome contact sites (Mattiazzi Ušaj et al., 2015). Furthermore, peroxisomes were found in close proximity to sites enriched in pyruvate dehydrogenase (PDH) complex responsible for acetyl-CoA synthesis in the mitochondrial matrix, further validating the organelles’ co-dependency in their metabolic processes (Cohen & Schuldiner, 2011; Mattiazzi Ušaj et al., 2015; Shai et al., 2016). In a high throughput screen, Fzo1 and Pex34 were found to contribute to the formation of mitochondria-peroxisomes tethers (Shai et al., 2016). In addition to their close physical and metabolic relationships, mitochondria and peroxisomes also utilize many similar outer membrane proteins, such as the same fission machinery, for their division.

A central component of this fission machinery is Fis1, a tail anchored (TA) protein that can be targeted to both mitochondrial and peroxisomal membranes in yeast, plants, and mammalian cells (Koch et al., 2005; Kuravi et al., 2006; M. Schrader et al., 2016). The adapter protein Mdv1, which, is recruited by Fis1 to the site of fission, engages in turn the dynamin-like protein Dnm1 that mediates the final fission step (Bleazard et al., 1999; Mozdy et al., 2000; Shaw & Nunnari, 2002; Yoon et al., 2003). Notably, mammalian homologues of these fission components were also found to be dually localized to mitochondria and peroxisomes. Combined defects in the organellar fission have been linked to several pathophysiological conditions and therefore it is important to understand the biogenesis of proteins involved in these cellular machineries (Ong et al., 2013; Schrader et al., 2022). Additional examples for proteins dually localized to both mitochondria and peroxisomes are the TA protein Gem1 (MIRO1 in mammals) and the ATPase Msp1 (ATAD1 in mammals)(Costello et al., 2017; Okreglak & Walter, 2014).

While a clear mechanism for targeting TA proteins to the secretory pathway has been worked out (Schuldiner et al., 2008; Stefanovic & Hegde, 2007), the mechanism involved in the correct targeting of TA proteins to mitochondria is still widely unknown. It has been previously shown that optimal hydrophobicity of the transmembrane domain and the presence of charged residues is essential for the correct targeting of TA proteins to mitochondria and/or peroxisomes (Bittner et al., 2022a; Borgese et al., 2007; Costello et al., 2017). While the mitochondrial import (MIM) complex was suggested to promote the biogenesis of the TA proteins Fis1 and Gem1 (Doan et al., 2020), other studies have also reported insertion of Fis1 to the mitochondrial outer membrane (MOM) and to lipid vesicles in an unassisted manner (Krumpe et al., 2012, Vitali et al., 2020).

For peroxisomal TA proteins, a dedicated pathway for membrane targeting – mediated by Pex19 and Pex3 exists (Fujiki et al., 2006). Nascent TA proteins with a peroxisomal membrane targeting signal (mPTS) are recognized by Pex19 in the cytoplasm and delivered to the peroxisomal membrane by interacting with the membrane receptor Pex3 (Chen et al., 2014; Götte et al., 1998). Surprisingly, depletion of *PEX19* resulted in reduced steady state levels of Fis1 and Gem1 also in mitochondrial fractions, suggesting an unexpected contribution of Pex19 to the biogenesis of mitochondrial TA proteins (Cichocki et al., 2018). Hence, despite some understanding of how TA proteins arrive to either mitochondria or peroxisomes, how the distribution of such dually localized proteins is regulated is still quite puzzling.

To obtain new insights on the mechanisms that control dual targeting of proteins to both mitochondria and peroxisomes and to unravel novel factors involved in this process, we used a high-throughput visual screen with fluorescently labeled TA proteins in the bakers’ yeast *Saccharomyces cerevisiae*. We found that the deletion of the uncharacterized gene *YNL144C* (in this study, re-named as Mitochondria and Peroxisomes Factor 1 (Mpf1)), as well as each of the paralogous proteins, *TOM70* and *TOM71* led to an enhanced localization of Fis1 to peroxisomes. Accordingly, overexpressing Tom71 caused Fis1 to localize more to mitochondria. We further characterized Mpf1 and identified it as an unstable protein on the surface of mitochondria controlled by the presence of Tom70 and Tom71. Collectively, our findings describe the involvement of Mpf1, Tom70, and Tom71 in regulating the dual distribution of Fis1 and Gem1 to mitochondria and peroxisomes.

## Results

### A high throughput visual screen reveals candidates that affect the dual distribution of TA proteins to mitochondria and peroxisomes

Some yeast TA proteins like Fis1 and Gem1 are found on both organelles - mitochondria and peroxisomes. To identify factors that influence the dual distribution of these proteins, we decided to employ a high throughput microscopy screen. To visualize the TA proteins of interest, we generated strains expressing mCherry tagged versions of either Fis1 or Gem1 on the background of a peroxisomal marker, Pex3-GFP. Then, using an automated procedure (Cohen & Schuldiner, 2011), we introduced these two tagged proteins into the background of a collection containing deletion strains of all the yeast non-essential genes and depletion strains of all the essential ones (Figure 1A). Thus, a new collection of deletion/depletion strains expressing the mCherry tagged TA protein along with GFP tagged Pex3 was created. This new collection was subjected to a high-throughput microscopy screen to identify those strains where the distribution of the TA proteins Fis1 and Gem1 between mitochondria and peroxisomes is altered (Figure 1A).

**Figure 1:**
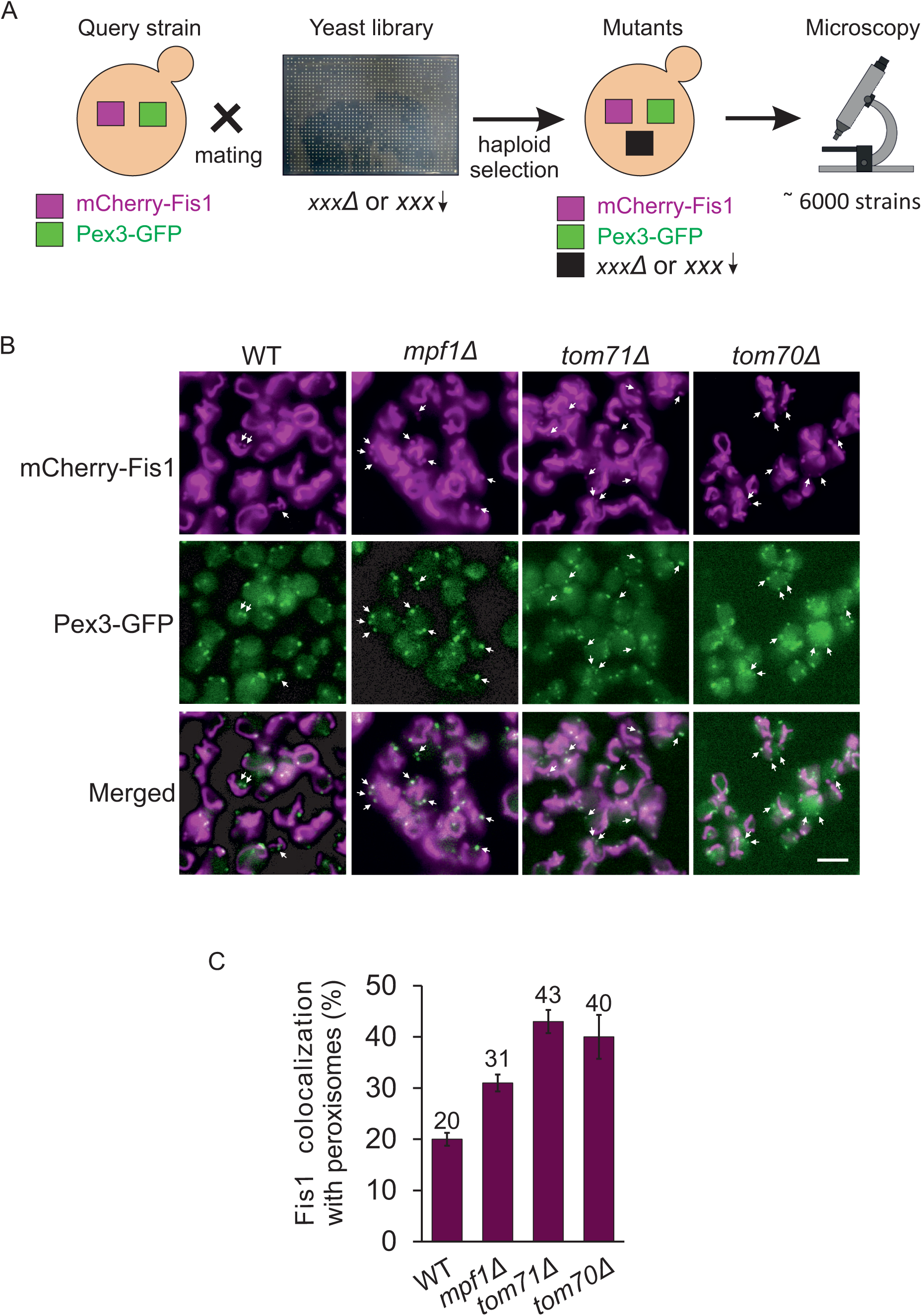
A high-throughput microscopy screen reveals proteins that affect dual distribution of Fis1. **(A)** Illustration of the screen aiming to find factors that affect dual targeting of Fis1. mCherry-Fis1 and Pex3-GFP were integrated into a yeast deletion and depletion libraries. The resultant strains, each containing a unique gene deletion/depletion and carrying the fluorescently labelled target proteins were visualized using automated microscopy. **(B)** Representative images of WT and three deletions strains with altered distribution of Fis1. The phenotype was observed by detecting co-localization of mCherry-Fis1 with Pex3-GFP (shown with white arrows). Scale bar, 5 µm. **(C)** Quantification of the co-localization of mCherry-Fis1 with peroxisomes. Total number of peroxisomes (visualized by Pex3-GFP) were counted in 100 cells in each of three independent experiments. Subsequently, the percentage of mCherry-Fis1 puncta co-localized with peroxisomes was determined. The graph represents the average of three independent experiments, error bars represent standard error.

While many proteins altered the distribution between the two organelles, it appeared that this was often secondary to biogenesis and/or morphology defects of the respective organelles. Hence, we decided to consider as a real hit only strains that fulfilled two criteria: (i) have normal biogenesis of peroxisomes (as reflected by the number of GFP puncta structures), and (ii) display normal mitochondrial morphology, as observed with the mCherry-Fis1. Considering these requirements, the screen allowed us to identify several proteins that might influence the distribution of Fis1 and Gem1 to peroxisomes (see Table S1 for the full list). Further manual examination of the complete list led us to focus on the uncharacterized protein Ynl144c (that we re-named as <u>m</u>itochondrial and <u>p</u>eroxisomal factor 1, Mpf1), and the two paralogue proteins Tom70 and Tom71. The absence of each of these three proteins led to a greater co-localization of mCherry-Fis1 with Pex3-GFP stained peroxisomes compared to control cells (Figure 1B and C). When we quantified this phenotype, we could detect co-localization of mCherry-Fis1 with 20% of peroxisomes in the wild type (WT) cells and this number was considerably increased in cells lacking Mpf1, Tom70, or Tom71 (Figure 1C).

To investigate whether this observation was limited only to mCherry-Fis1, we quantified the co-localization of mCherry-Gem1 with Pex3-GFP and observed the same trend, indicating that the identified proteins have a general effect on dually distributed TA proteins (Figure S1). We further confirmed that these observations were not the result of fragmented mitochondria that were misinterpreted for peroxisomal puncta. To this aim, we visualized mitochondrial morphology by staining the organelle with Om45-GFP and observed that the mitochondrial morphology was not altered in *mpf1*Δ and *tom71*Δ cells as compared to control cells (*tim13*Δ) (Figure S2). We noticed a slightly altered mitochondrial morphology in *tom70*Δ cells (Figure S2), which is not surprising considering the functions of Tom70 as an important mitochondrial import receptor and a docking site for cytosolic chaperones (Backes et al., 2021; Kreimendahl & Rassow, 2020; Yamamoto et al., 2009; Young et al., 2003). Collectively, the visual screen identified Mpf1, Tom70, and Tom71 as potential factors affecting the dual distribution of TA proteins.

### Physical separation of mitochondria and peroxisomes validates the involvement of the identified hits in the dual distribution of Fis1

To confirm by another unrelated approach, that the candidates that we picked by the visual screen truly affect the dual distribution of Fis1 to mitochondria and peroxisomes, we monitored the distribution of Fis1 by subcellular fractionation. To obtain optimal separation, the cells were grown on oleate as a carbon source, a condition known to induce proliferation of peroxisomes. After obtaining a crude mitochondrial fraction, we used centrifugation of a Histodenz gradient to separate mitochondria from peroxisomes and 12 fractions of the gradient were collected. Our protocol could nicely differentiate between the two organelles despite their strong physical associations: Tom20, a *bona-fide* mitochondrial marker protein was enriched in the first four fractions (lanes 1-4), whereas the peroxisomal marker protein Pex14 was found preferentially in the last four fractions (9-12). As expected for a dually targeted protein, Fis1 was present in both sets of fractions (lanes 1-4 and 9-12) (Figure 2A). Fis1 levels in the various fractions were quantified and the total amount in fractions 1-4 was considered as mitochondrial Fis1 whereas the material in fractions 9-12 was counted as peroxisomal population.

**Figure 2:**
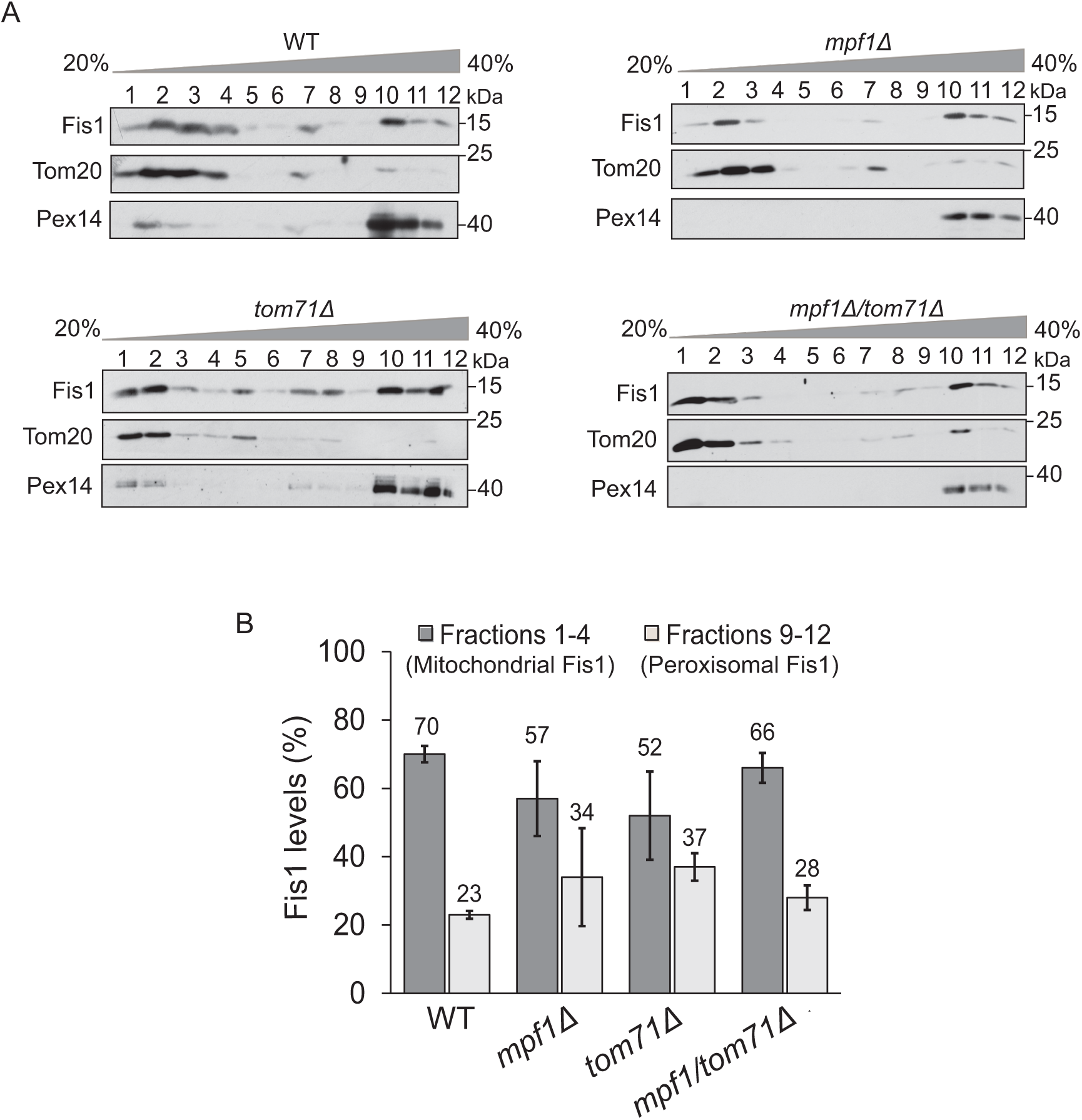
Physical separation of mitochondria and peroxisomes validates the hits. **(A)** Gradient centrifugation procedure was employed to separate mitochondria and peroxisomes from the indicated strains and 12 fractions from the top of the gradient were collected. The fractions were analyzed by SDS-PAGE and immunodecoration with antibodies against Fis1 (dually localized to mitochondria and peroxisomes), Tom20 (mitochondrial marker), and Pex14 (peroxisome marker). **(B)** The intensities of Fis1 obtained in each fraction was quantified and the sum of all the 12 intensities was set to 100%. Fis1 signal in fractions 1-4 was considered mitochondrial, while that within fractions 9-12 was designated as peroxisomal. The graph represents the average of three independent experiments, error bars representing standard error.

Using this approach, we compared the distribution of native Fis1 in wild type (WT) cells to that in the mutated strains. Whereas in WT cells, 70% of Fis1 were found to be mitochondrial, this portion decreased to 57% and 52% in *mpf1*Δ and *tom71*Δ cells, respectively (Figure 2A and B). In parallel to the decrease in mitochondrial Fis1, we observed an increase in the peroxisomal portion of the protein. We found 23% of Fis1 in peroxisomes in WT cells whereas this fraction had increased to 34% and 37% in *mpf1*Δ and *tom71*Δ cells, respectively (Figure 2A and B). Unfortunately, we could not analyze the distribution of Fis1 in *tom70Δ* cells because, for unknown reasons, the separation of mitochondria from peroxisomes did not work well with cells from this strain. Hence, this technique could not validate Tom70 as a factor regulating the distribution of TA proteins.

We were then interested to explore the distribution phenotype upon the parallel deletion of both *MPF1* and *TOM71*. Surprisingly, we observed that in *tom71Δ/mpf1*Δ double deletion cells, the alteration in the Fis1 distribution was less profound than in the corresponding single deletion strains (Figure 2A and B). It might be that loss of both Mpf1 and Tom71 leads to compensatory upregulation of alternative factors to restore mitochondrial Fis1 levels to those observed in control cells. Taken together, the physical separation of mitochondria from peroxisomes confirmed the (direct or indirect) involvement of Mpf1 and Tom71 in regulating the dual distribution of Fis1 between mitochondria and peroxisomes.

### Tom71 has a unique role in Fis1 distribution, setting it apart from Tom70

Tom71 is a paralogue of Tom70, sharing 53% sequence identity, whose abundance is rather low – only about 10% of the levels of Tom70 (Morgenstern et al., 2021; Schlossmann et al., 1996). Tom70 plays a pivotal role as a mitochondrial import receptor and chaperones’ docking site and is required for the biogenesis of many mitochondrial proteins (Backes et al., 2021; Kreimendahl & Rassow, 2020; Yamamoto et al., 2009; Young et al., 2003). Tom70 and Tom71 are thought to have overlapping functions, and a specialized role unique to Tom71 has not been reported yet.

Our work suggested a unique role for Tom71 since its deletion led to a distribution phenotype even in the presence of Tom70. To better understand the involvement of both proteins in the dual distribution of Fis1, we created strains where each of the protein is over-expressed. To that aim, the endogenous promoter of *TOM70* in WT, *mpf1*Δ and *tom71*Δ cells was replaced by the strong *GPD* promoter, resulting in a dramatic overexpression (Figure 3A). We then separated mitochondria and peroxisomes and quantified mitochondrial and peroxisomal levels of Fis1 (Figure 3B and C). We observed that elevated levels of Tom70 led to a correction of Fis1 distribution in *mpf1*Δ cells with 70% mitochondrial and 25% peroxisomal Fis1, similar to the distribution observed in WT cells (Figure 3B and C). Thus, it seems that the role of Mpf1 in regulating Fis1 distribution is dispensable in the presence of higher amounts of Tom70. Interestingly, only a partial correction of Fis1 distribution was observed upon overexpression of Tom70 in *tom71*Δ cells with 64% in mitochondria and 30% in peroxisomes (Figure 3B and C), indicating that Tom71 has a unique role in Fis1 distribution that cannot be replaced by elevated levels of Tom70.

**Figure 3:**
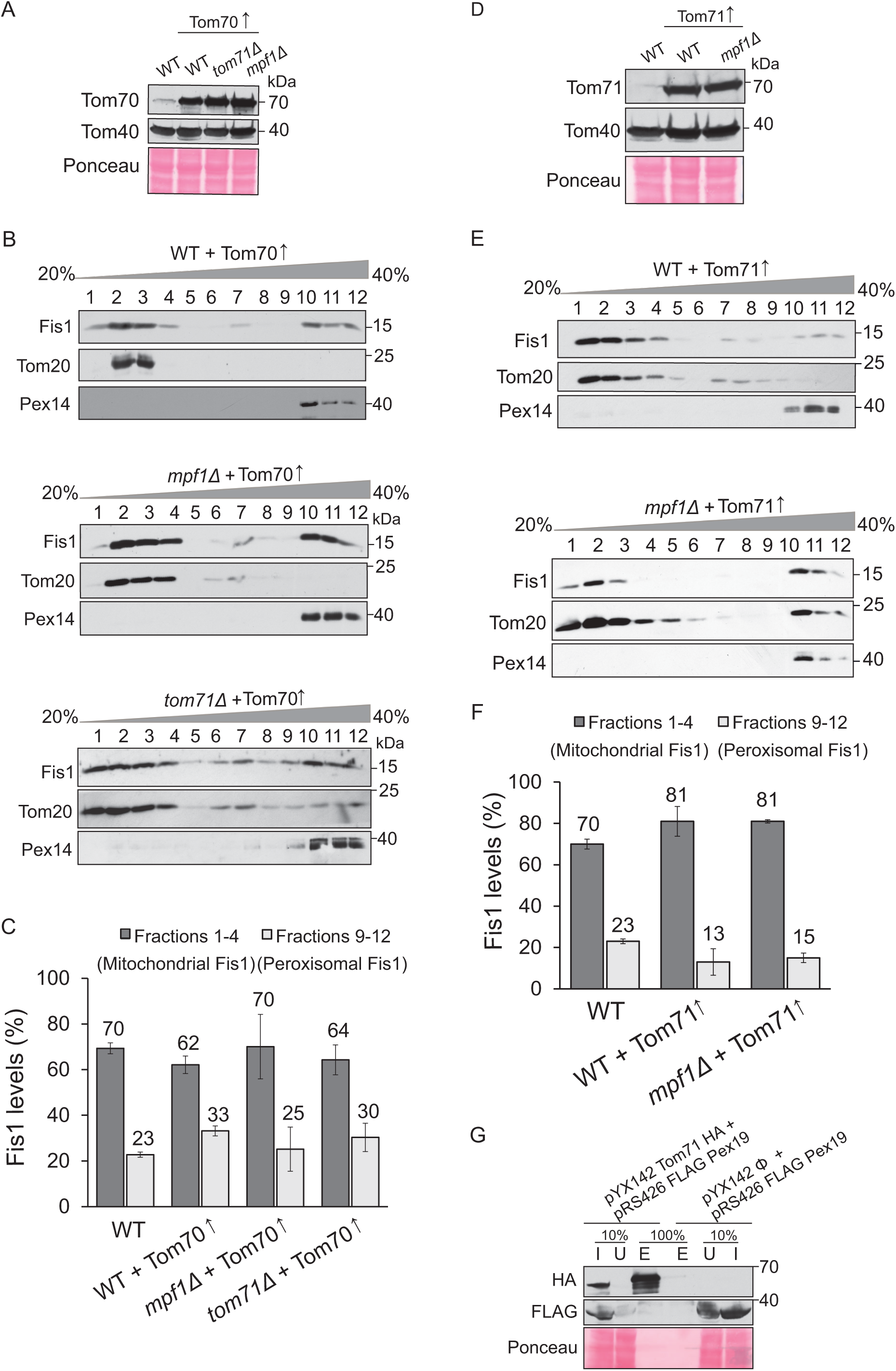
Tom71 has a unique effect on the distribution of Fis1. **(A)** Tom70 was overexpressed in the indicated strains by replacing the endogenous promoter with the *GPD* promoter. Cells of the resulting strains were grown on galactose and whole cell lysate was obtained by alkaline lysis. Extracted proteins were analyzed by SDS-PAGE and immunodecoration with the indicated antibodies. Ponceau staining was employed to verify equal loading in all lanes. **(B, C)** Gradient separation of mitochondria and peroxisomes from the indicated strains were performed as described in the legend to Fig. 2A and B. **(D)** Tom71 was overexpressed in the indicated strains by replacing the endogenous promoter with the *GPD* promoter. Proteins from the obtained strains were analyzed as described in part (A). **(E-F)** Gradient separation of mitochondria and peroxisomes from the indicated strains were performed as described in the legend to Fig. 2A and B. Note: to allow easier comparison, the Fis1 levels in WT cells in panels C and F were taken from Figure 2B. The graph represents the average of three independent experiments, error bars representing standard error. **(G)** Cells expressing either Flag-Pex19 alone or co-expressing Flag-Pex19 and Tom71-HA were lysed with Triton X-100 and the suspension was incubated with anti-HA beads. Fractions representing the input (I), unbound material (U), and the eluate (E) were analyzed by SDS-PAGE and immunodecoration with the indicated antibodies.

Remarkably, although we expected that overexpression of Tom70 in WT cells would drive Fis1 distribution more towards mitochondria, we observed a minor reduction in mitochondrial Fis1 (62%) and a slight increase in peroxisomal Fis1 (33%) (Figure 3B and C). This observation is specially intriguing considering the report that the biogenesis of many mitochondrial proteins is enhanced upon overexpression of Tom70 (Liu et al., 2022). We can speculate that targeting of Fis1 towards mitochondria might prefer Tom71 over Tom70 and over-crowding the mitochondrial surface with Tom70 and/or engaging Tom71 in Tom71/Tom70 heterodimers creates a competing effect, thereby reducing Fis1 levels in mitochondria.

To further study the role of Tom71 in regulating the distribution of Fis1, we next constructed WT and *mpf1*Δ strains where Tom71 expression is under the control of the strong *GPD* promoter and could confirm the massive overexpression of Tom71 in these cells (Figure 3D). Then, we separated mitochondria and peroxisomes from these strains and quantified the levels of mitochondrial and peroxisomal Fis1 (Figure 3E and F). Interestingly, we observed that Tom71 over expression in both WT and *mpf1*Δ cells led to an increased distribution of Fis1 towards mitochondria (Figure 3E and F). These findings substantiate the independent and unique contribution of Tom71 to the targeting of Fis1 to mitochondria.

It has previously been shown that the cytosolic chaperone/receptor Pex19 assists the biogenesis of mitochondrial Fis1 (Cichocki et al., 2018). Since overexpression of Tom71 drove 81% of Fis1 towards mitochondria, we wondered whether Tom71 functions as a receptor for Pex19 on the surface of mitochondria. To test this possibility, we created a strain where Tom71-HA and Flag-Pex19 are co-overexpressed in *tom70*Δ cells. We deleted Tom70 from these cells to prevent potential competition of Tom70 in binding to Pex19. Next, pull down analysis to detect potential interaction was performed. However, we could not co-elute Flag-Pex19 with Tom71-HA (Figure 3G). This outcome proposes that the proteins might not interact, or that the interaction is very transient. Taken together, these results demonstrate a unique role of Tom71, whose absence cannot be compensated by Tom70, while the contribution of Mpf1 to regulating Fis1 distribution is dispensable upon overexpression of either Tom70 or Tom71.

### Deletion of *MPF1* is beneficial to cells grown on oleic acid

Since Mpf1 is an uncharacterized protein with an unknown function, we wanted to investigate whether loss of Mpf1 would have an effect on growth of yeast cells. We noticed that *mpf1*Δ cells grew similar to WT cells when glucose (YPD), glycerol (YPG), or oleate (YPO) were used as carbon sources in a rich medium. However, when oleic acid was used as the sole carbon source on a synthetic medium, cells lacking Mpf1 grew better than WT cells (Figure 4).

**Figure 4:**
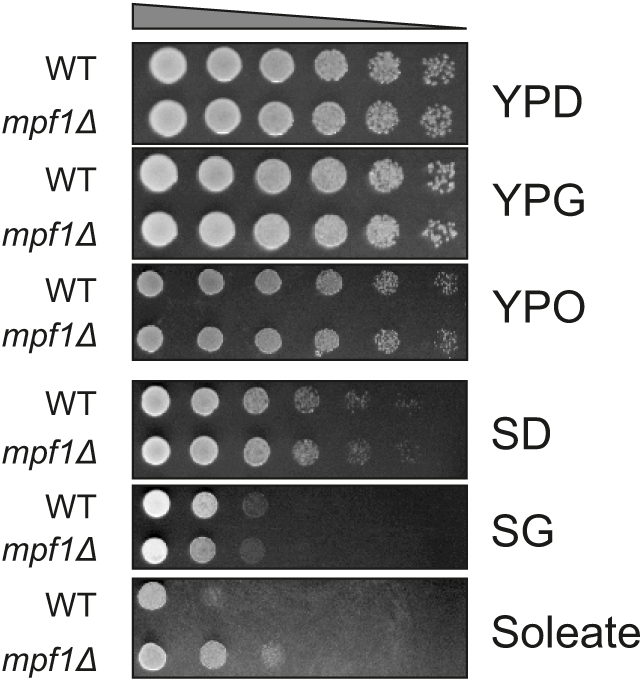
Loss of Mpf1 is beneficial for cells grown on oleic acid as a carbon source. Growth of wild-type (WT) and *mpf1Δ* cells at 30°C was analyzed by drop-dilution assay. The cells were grown on either rich media (YP) or synthetic media (S) containing glucose (YPD or SD), glycerol (YPG or SG), or oleate (YPO or SOleate).

In yeast, β-oxidation of fatty acids such as oleate takes place solely in peroxisomes (Hiltunen et al., 2003). Thus, yeast cells require fully functional peroxisomes for optimal growth on oleate as the exclusive carbon source. Previous studies have shown that absence of Fis1 reduces the number of peroxisomes in cells grown on oleate (Kuravi et al., 2006). Moreover, re-directing Fis1 only to peroxisomes by expressing Fis1-Pex15 fusion protein increased Dnm1-dependent peroxisome fission, and thereby increased the number of peroxisomes per cell (Motley et al., 2008). Hence, the better growth of *mpf1*Δ cells on oleate can be explained by the increased portion of Fis1 in peroxisomes in these cells which in turn, enhances the number of peroxisomes and thus improves the utilization of oleate.

### Mpf1 is a highly unstable protein

A previous high throughput study proposed that Mpf1 might be a substrate of Grr1, an SCF ubiquitin ligase complex subunit. Both Mpf1 and its uncharacterized paralog, Yhr131c were suggested to interact with Grr1 and were reported to be partially stabilized in *grr1*Δ cells (Mark et al., 2014). To verify this previous report, we transformed a plasmid encoding Mpf1-3HA into WT and *grr1*Δ cells and monitored the life span of Mpf1 in these cells. In line with the previous findings, we observed that Mpf1 is indeed a highly unstable protein in WT cells and is almost completely degraded within 45 minutes after inhibition of translation by addition of cycloheximide (CHX) (Figure 5A and B). However, in our hands, the deletion of *GRR1* had only a minor effect on the stability of Mpf1 (Figure 5 A and B), suggesting that there are other factors that affect the lifespan of this protein.

**Figure 5:**
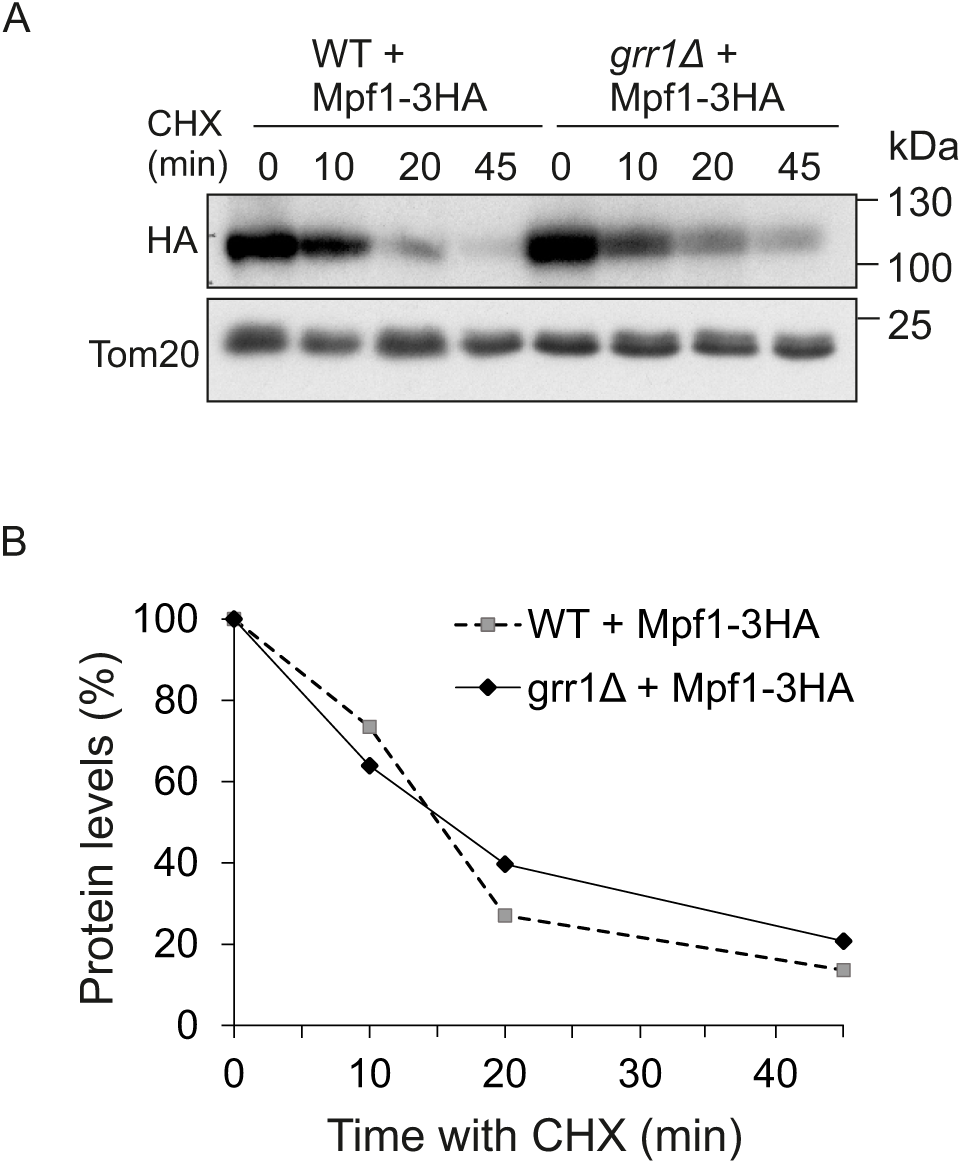
Mpf1 is an unstable protein. **(A)** WT and *grr1Δ* cells were transformed with a vector encoding Mpf1-3HA. The cells were grown on SD-Ura and then at time = 0 the translation inhibitor cycloheximide (CHX) was added. Cells were further incubated and proteins were extracted at each time point by alkaline lysis and analyzed by SDS-PAGE and immunodecoration with antibodies against HA or Tom20 (as a loading control). **(B)** The bands representing either Mpf1-3HA or Tom20 were quantified and for each lane, the intensity of the band corresponding to Mpf1-3HA was normalized to the loading control (Tom20). The signal at time point = 0 was set to 100%. One representative experiment out of three independent ones is shown.

### Mpf1 loosely associates with the mitochondrial outer membrane

To further characterize Mpf1, we aimed to determine its subcellular location. WT cells expressing Mpf1-3HA were fractionated into whole cell lysate (WCL), ER, cytosol and mitochondria and these fractions were analyzed by Western blotting. Antibodies recognizing marker proteins for the mitochondria (Tom40), ER (Erv2), cytosol (Hexokinase) and peroxisomes (Pex14) were used to verify the fractionation. We detected Mpf1-3HA mainly in the mitochondrial fraction with some portion also in the ER fraction. Given the presence of the peroxisomal marker Pex14 in both fractions, we could only conclude that Mpf1-3HA might localize to mitochondria, peroxisomes, and/or the ER (Figure 6A).

**Figure 6:**
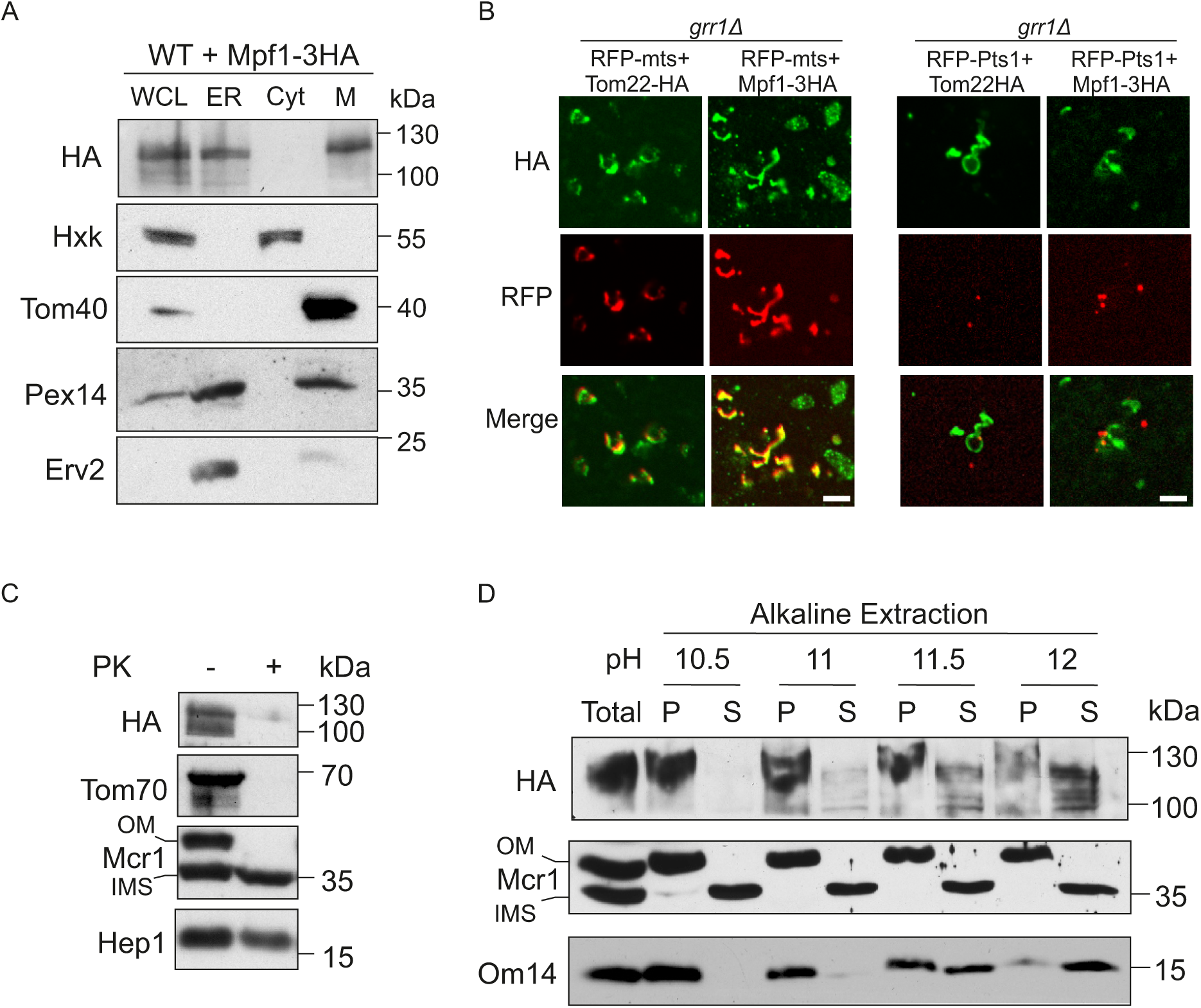
Mpf1 shows a loose association with the mitochondrial outer membrane. **(A)** Cells overexpressing Mpf1-3HA were subjected to subcellular fractionation. The isolated fractions of whole cell lysate (WCL), microsomes (ER), cytosol (Cyt), and mitochondria (M) were analyzed by SDS-PAGE and immunodecoration with the indicated antibodies. Tom40 (mitochondria), Hexokinase (cytosol), Pex14 (peroxisomes), and Erv2 (ER) were used as marker proteins. **(B)** Cells expressing Mpf1-3HA were analyzed by immunofluorescence microscopy. *grr1Δ* cells were used to increase the half-life of the protein. The HA tagged proteins were visualized with anti-HA antibody conjugated with Alexa Fluor™ 488. Tom22-HA, a *bona-fide* mitochondrial protein was used as a control for the procedure. To visualize mitochondria and peroxisomes, the cells expressing the HA-tagged proteins were co-transformed with MTS-RFP (mitochondrial targeting signal) or RFP-PTS1 (peroxisomal targeting signal 1). Scale bar, 5 µm. **(C)** Isolated mitochondria from cells expressing Mpf-3HA were either left intact (-PK) or treated with proteinase K (+PK). Then, the samples were analyzed by SDS-PAGE and immunodecoration with the indicated antibodies. Tom70 and Mcr1_OM_ are exposed on the mitochondrial surface whereas Mcr1_IMS_ and Hep1 (matrix) are protected by mitochondrial membranes. **(D)** Isolated mitochondria from cells expressing Mpf-3HA were subjected to alkaline extraction using solution at the indicated pH values. “Total” represents untreated mitochondria. Membrane proteins were isolated in the pellet (P) fraction and soluble and membrane-peripheral proteins in the supernatant (S) fraction. The samples were analyzed by SDS-PAGE and immune-decoration against the specified antibodies. Mcr1_OM_ and Mcr1_IMS_ served as controls for integral membrane protein and soluble protein, respectively. Om14 acted as a control for MOM-associated protein extractable under extreme alkaline conditions.

To obtain a more precise localization of Mpf1, we then chose to employ fluorescence microscopy techniques. Tagging Mpf1 with GFP led to a cytosolic staining, which was not in line with the subcellular fractionation assays. This observation can be explained by either cleaving of the GFP tag and/or mis-targeting due to the bulky GFP moiety. Hence, we opted for immunofluorescence (IF) assays. Anti-HA antibodies conjugated to a fluorophore were used to detect Mpf1-3HA. As a control for the IF technique, we also visualized Tom22-HA, a *bona-fide* mitochondrial outer membrane protein. We observed that Mpf1-3HA stained tubular structures that co-localize with RFP fused to mitochondrial targeting signal (RFP-MTS) but not with RFP-PTS1 (Peroxisomal targeting signal 1) (Figure 6B). These findings support the notion that Mpf1-3HA localizes to mitochondria.

Since mitochondria have four different sub-compartments, we investigated the sub-mitochondrial localization of Mpf1-3HA by treating mitochondria isolated from cells expressing Mpf1-3HA with proteinase K (PK). We found Mpf1-3HA to be susceptible to PK digestion comparable to that of the surface proteins Tom70 and an Mcr1 isoform on the outer membrane (Mcr1_OM_) (Figure 6C). As expected for mitochondrial internal proteins, the Mcr1 isoform in the intramembrane space (Mcr1_IMS_) and the matrix protein Hep1 were protected from PK by the outer membrane and both outer and inner membranes, respectively (Figure 6C). Hence, we concluded that Mpf1 is exposed to the cytosol on the surface of mitochondria.

We next determined whether Mpf1-3HA is an integral or peripheral membrane protein by performing alkaline extraction of mitochondrial proteins followed by centrifugation to separate membrane embedded proteins in the pellet from soluble and peripheral membrane proteins in the supernatant. To obtain a better resolution of the assay, we performed it under varying pH conditions. The alkaline pH decreases non-covalent protein-protein interactions and releases peripheral membrane proteins to the supernatant (Kim et al., 2015). As expected, Mcr1_OM_ isoform remained in the pellet fractions under all pH conditions, confirming its behavior as a *bona fide* integral membrane protein. In contrast and as anticipated, the soluble IMS isoform of Mcr1 was in the supernatant fraction under all the employed conditions (Figure 6D). We found that Mpf1-3HA was in the membrane fraction only in milder extraction condition (pH 10.5). As the pH was raised to 11, 11.5, and 12, increasing amounts of Mpf1 were found in the supernatant fraction. This behavior is similar to that of the outer membrane proteins Om14, which is known to be partially extractable under alkaline conditions (Burri et al., 2006; Zhou et al., 2022) (Figure 6D). Altogether, these results indicate that Mpf1-3HA is peripherally associated with the cytosolic face of the mitochondrial OM.

### The combined loss of Tom70 and Tom71 affects the steady state levels of Mpf1

Considering the location of Mpf1 on the mitochondrial surface, we wondered whether Tom70, Tom71, or both are involved in targeting of Mpf1 to the organelle. Several studies on Tom70 and its paralog Tom71 (to a lesser extent) have identified a tetratricopeptide (TPR) structure that can bind to the cytosolic chaperones Hsp70 and Hsp90 and enable the recruitment of several precursor proteins together with chaperones to the mitochondrial surface (Backes et al., 2018; Jores et al., 2018; Young et al., 2003; Zanphorlin et al., 2016). Moreover, microscopy analysis of many GFP tagged proteins showed their reduced levels upon the absence of Tom70/71 (Backes et al., 2021). To investigate potential involvement of Tom70/71 in the biogenesis of Mpf1, we monitored the steady state levels of Mpf1-3HA in *tom70*Δ, *tom71Δ,* and double deletion *tom70Δ/71*Δ cells. Of note, in the absence of either Tom70 or Tom71 alone, Mpf1-3HA levels were comparable to those in WT cells. However, double deletion of both proteins resulted in tenfold decrease compared to WT cells (Figure 7A and B).

**Figure 7:**
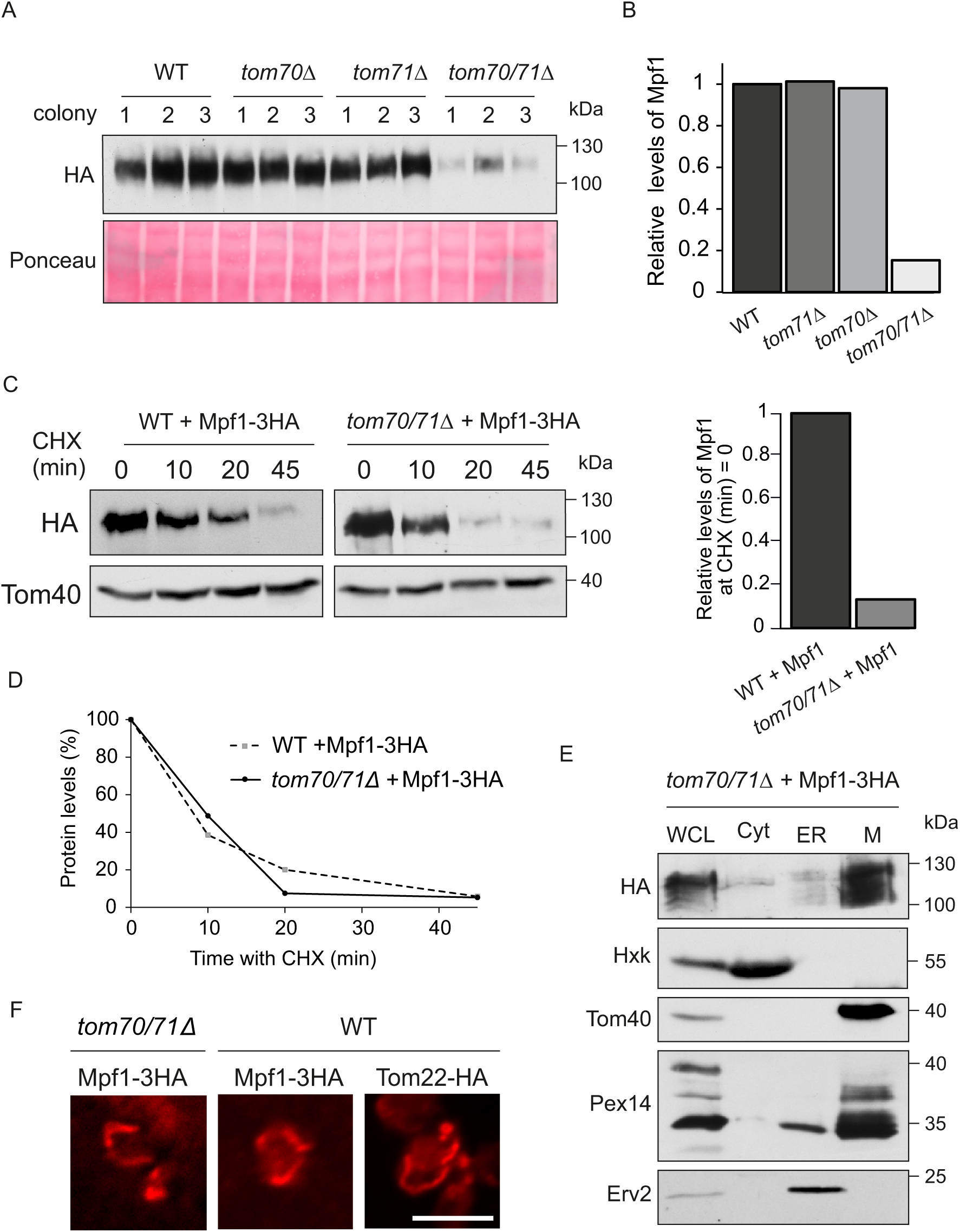
The absence of Tom70 and Tom71 reduces the expression of Mpf1. **(A)** Proteins were extracted from the indicated cells (three independent colonies) expressing Mpf1-3HA. Samples were analyzed by immunodecoration with HA antibody. Ponceau staining was used as the loading control. **(B)** The bands representing Mpf1-3HA were quantified and for each lane normalized to the intensity of the Ponceau staining. The average of the three colonies for each strain was calculated and the value for WT cells was set as 1. **(C)** Left panel: WT and *tom70/71Δ* cells expressing Mpf1-3HA were subjected to cycloheximide (CHX) assay as described in the legend to Fig. 5A. Right panel: Quantification of Mpf1-HA levels relative to Ponceau at time point = 0. **(D)** The bands corresponding to Mpf1-3HA in the experiment presented in panel (C) were quantified as described in the legend to Fig. 5B. One representative experiment out of three independent ones is presented. **(E)** Subcellular fractionation of *tom70/71Δ* cells expressing Mpf1-3HA. The isolated fractions of whole cell lysate (WCL), microsomes (ER), cytosol (Cyt), and mitochondria (M) were analyzed by SDS-PAGE and immunodecoration with the indicated antibodies. Tom40 (mitochondria), Hexokinase (cytosol), Pex14 (peroxisomes), and Erv2 (ER) were used as marker proteins. **(F)** Immunofluorescence microscopy to visualize Mpf1-3HA in *tom70/71Δ* and WT cells. Tom22-HA was used as a control for the IF procedure. The HA-tagged proteins were visualized using an anti-HA antibody conjugated with Alexa Fluor™ 594. Scale bar, 5 µm.

To understand the reason for such a dramatic reduction in the steady-state levels, we investigated whether the stability of Mpf1-3HA was affected in *tom70Δ/71*Δ cells. Though the relative levels of Mpf1-3HA was tenfold lower in the double deletion cells at the beginning of the assay (time-point 0), the life span of the protein in the mutated cells was comparable to that in WT cells (Figure 7C and D). Furthermore, both subcellular fractionation and immunofluorescence microscopy revealed that Mpf1-3HA still localized predominantly to the mitochondria in *tom70Δ/71*Δ cells (Figure 7E and F).

Recent studies suggested that Tom70 can be involved in signaling through multiple transcription factors to control the transcription levels of genes encoding for many mitochondrial proteins. Accordingly, upon the selective removal of Tom70 from the mitochondrial surface, the levels of mRNAs encoding mitochondrial proteins were reduced (Liu et al., 2022). Along the same line, our RT-qPCR analysis revealed that the transcript levels of endogenous *MPF1* as well as overexpressed *MPF1-3HA* were reduced by around 50% as compared to WT cells upon the deletion of both *TOM70/71* (Figure S3). These lower mRNA levels can explain (at least partially) the dramatically reduced levels of Mpf1 protein in the double deletion strain. It should be mentioned that such reduction in the detection of mRNA can result from less transcription of *MPF1,* increased degradation of the mRNA, and/or sequestering of the mRNA to P-bodies. Collectively, our findings suggest an involvement of both Tom70 and Tom71 in the transcriptional control of Mpf1-3HA. Although the mRNA and the protein steady-state levels of Mpf1-3HA are dramatically reduced in the absence of Tom70/71, Mpf1-3HA still localizes to mitochondria, suggesting the involvement of other factor(s) in its biogenesis.

### The PH domain of Mpf1 is involved in the stability of the protein but not in its localization

Large scale studies and structural analysis predicted the presence of a Pleckstrin homology (PH) domain in Mpf1 (Gallego et al., 2010; Isakoff et al., 1998; Lemmon, 2004). This domain can interact with phosphoinositide (PI) species as well as other lipids on biological membranes. Previous attempts to investigate binding of a recombinant PH domain from Mpf1 to PI were unsuccessful due to inadequate quantities (Yu et al., 2004). PH domains occur in a wide range of proteins with varying functions, and are stretches of ∼120 amino acid residues with two anti-parallel β-sheets followed by a C-terminal α-helix (Riddihough, 1994). Membrane targeting of PH domains that strongly bind to PIs could be abolished by mutating the basic residues in the β1/β2 loop (Yu et al., 2004). To investigate if the PH domain of Mpf1 is essential for the stability and/or subcellular localization of the protein, we created a mutant of Mpf1 with lysine and arginine residues in the PH domain replaced by alanine (K144A, K147A, and R157A) (Figure 8A, Mpf1(PH*)-3HA). Compared to regular Mpf1-3HA, Mpf1(PH*)-3HA exhibited elevated stability, with steady-state levels ∼60% higher than that of the native protein (Figure 8B and C). The PH domain is known to serve as a platform for protein-protein interactions (Lemmon, 2004; Scheffzek & Welti, 2012), and mutating the basic residues in the PH domain of Mpf1 might have disrupted its interactions with some factors of the proteasome degradation pathway and thereby increasing its stability. Of note, cells overexpressing Mpf1(PH*)-3HA grew slightly better than WT cells on oleate-containing medium (Figure 8D). It can be speculated that this variant might have a dominant negative effect on the recruitment of Fis1 to mitochondria and thus enhances the number of peroxisomes under these conditions.

**Figure 8:**
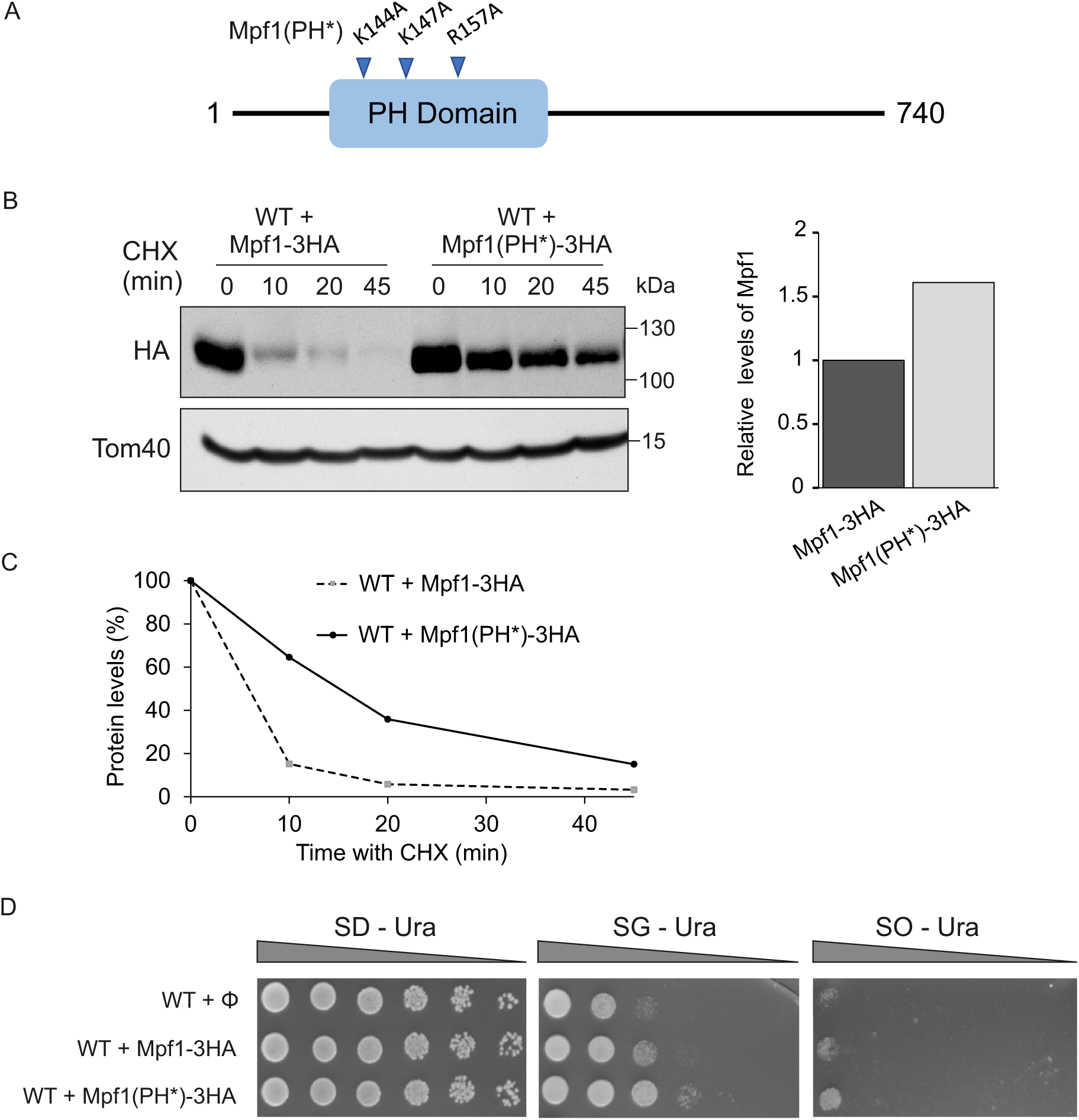
Mutating the Pleckstrin homology (PH) domain of Mpf1 stabilizes the protein. **(A)** Schematic diagram of the mutations in the PH domain of Mpf1 with K144, K147, and R157 replaced by alanine (A) residues (the mutant is indicated as Mpf1(PH*). The mutated basic residues in the in the β1/β2 loop of the PH domain are indicated with blue arrowheads. **(B)** Left panel: WT cells overexpressing either Mpf1-3HA or Mpf1(PH)*-3HA were subjected to cycloheximide (CHX) assay for the indicated time periods. Proteins were then extracted and analyzed by SDS-PAGE and immunodecoration with antibodies against either HA or Tom40 (as loading control). Right panel: Quantification of Mpf1-HA and Mpf1(PH)*-3HA levels relative to Tom40 at time point = 0. **(C)** Quantification of Mpf1-3HA and Mpf1(PH)*-3HA was performed as described in the legend to Fig. 5B. One representative experiment out of three independent ones is presented. **(D)** Growth analysis by drop dilution assay of WT cells harboring either an empty vector (Φ), a plasmid encoding Mpf1-3HA, or a plasmid encoding Mpf1(PH*)-3HA. Cells were grown at 30°C on synthetic media containing glucose (SD-Ura), glycerol (SG-Ura), or oleic acid (SO-Ura).

We next asked whether the basic residues in the PH domain of Mpf1 are required for its targeting to mitochondria and its association with the OM. Employing immunofluorescence microscopy and subcellular fractionation, we found that Mpf1(PH*)-3HA still localizes to the mitochondria (Figure 9A and B). Molecular modelling revealed that the PH domain of Mpf1 has an overall weak or no positive charge, and membrane targeting or binding to PIs may depend on strong positive charges (Yu et al., 2004). Furthermore, not all PH domains exhibit strong and specific interactions with PIs and many of them bind to PIs with low affinity and specificity. Effective binding of such “weak” PH domains to biological membranes might be strengthened by interactions with other membrane-bound proteins (Maffucci & Falasca, 2001). To explore this option, we then performed an alkaline extraction to see if the association of Mpf1(PH*)-3HA with the MOM is affected. Mpf1(PH*)-3HA predominantly remained in the pellet fraction in pH 10.5 and there was an increase of the protein portion in the supernatant only at pH 12, indicating that Mpf1(PH*)-3HA exhibits a similar, or even somewhat stronger association to the MOM than its native counterpart (Figure 9C). Collectively, these results suggest that mutating key residues in the PH domain of Mpf1 did not change its association with mitochondria.

**Figure 9:**
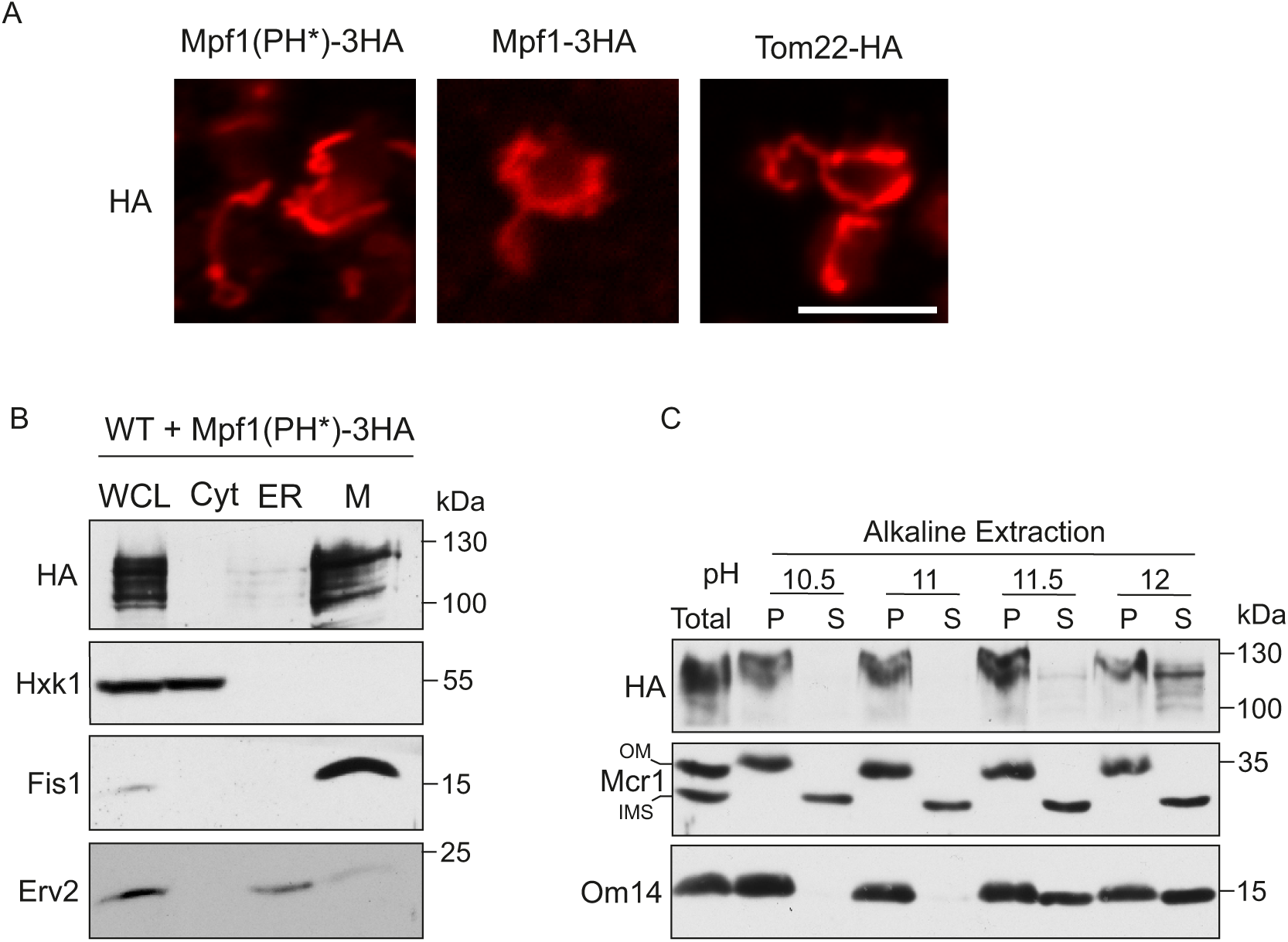
The mutations in the PH domain do not affect the location of Mpf1. **(A)** Immunofluorescence microscopy localization of Mpf1(PH*)-3HA in WT cells. Native Mpf1-3HA and Tom22-HA were used as a control for the procedure. The HA-tagged proteins were visualized using an anti-HA antibody conjugated with Alexa Fluor™ 594. Scale bar, 5 µm. **(B)** Subcellular fractionation of WT cells expressing Mpf1(PH*)-3HA. Cells were analyzed as described in the legend to Fig. 6A. (C) Isolated mitochondria from cells expressing Mpf1(PH*)-3HA were subjected to alkaline extraction as described in the legend to Fig. 6D.

## Discussion

Considering the number of shared proteins between mitochondria and peroxisomes and the *de novo* formation of peroxisomes via mitochondria-derived vesicles (at least in mammalian cells), several studies speculate that peroxisomes evolved to facilitate the quality control of mitochondria under periods of stress and to relieve mitochondria from the burden of hosting oxidation enzymes of the β-oxidation pathway (Bittner et al., 2022b; Speijer, 2017). This symbiotic relationship between mitochondria and peroxisomes might have allowed peroxisomes to utilize several mitochondrial proteins for their own needs like those proteins involved in fission (Fis1) and quality control (Msp1). Moreover, mitochondria contribute indirectly to the targeting of the phosphatase Ptc5p to peroxisomes via a mitochondrial transit. These cross-talks are proposed to eventually lead to targeting of these proteins to both organelles when peroxisomes became autonomous over time (Bittner et al., 2022b; Stehlik et al., 2020). Distribution of the dually localized TA protein Fis1 to mitochondria and peroxisomes is aided by Pex19 (Cichocki et al., 2018). However, regulatory mechanisms to understand how the distribution of these cytosolically synthesized TA proteins is controlled have remained elusive.

In this study, we used comprehensive techniques to find factors that regulate the dual distribution of TA proteins Fis1 and Gem1 to the membranes of mitochondria and peroxisomes. Using high-throughput microscopy screens, we first identified the involvement of Tom70, Tom71, and an uncharacterized protein Ynl144c (re-named as Mpf1 in this study) in regulating the distribution of Fis1 and Gem1 to mitochondria and peroxisomes. Subsequently, we verified the involvement of these candidates via subcellular fractionation assays and confirmed that in *tom71*Δ and *mpf1*Δ cells, Fis1 distributed more to peroxisomes. Surprisingly, the double deletion cells *tom71/mpf1*Δ showed Fis1 distribution comparable to WT cells. Since mitochondrial fission is crucial for the maintenance of healthy cells and the depletion of Fis1 leads to hyperfused mitochondria (Das & Chakrabarti, 2020; Hoppins et al., 2007), *tom71/mpf1*Δ potentially leads to activation of other factors to maintain proper distribution of Fis1 molecules. Further investigations on which factors could be upregulated in *tom71/mpf1*Δ cells would enhance our understanding of this regulatory mechanism. Unfortunately, due to technical difficulties, we were unable to optimally separate mitochondria and peroxisomes in *tom70*Δ and *tom70/71*Δ cells, and thereby the impact of the absence of Tom70 on the distribution of Fis1 could not be verified and remains unclear.

However, we were able to show that enhanced levels of Tom70 in *mpf1Δ* cells could fully correct the Fis1 distribution to WT levels but only partially reverse the distribution in *tom71Δ*. This observation suggests that Tom71 might have a more dominant role in affecting Fis1 distribution, that is not entirely rectified by higher levels of Tom70. Tom71 is a barely expressed paralog of Tom70 and until this study, a distinct function of Tom71 distinguishing it from Tom70 was not found. Our hypothesis that Tom71 plays a unique role in regulating Fis1 distribution is further validated by our finding that overexpression of Tom71 has a significant effect (compared to that of Tom70) in driving Fis1 distribution more towards mitochondria. A possible explanation for the effect of Tom71 is a putative role as a mitochondrial receptor for Pex19. However, so far, we could not detect a physical interaction between Tom71 and Pex19. It could be that the interaction is rather transient and might be dependent on the presence of substrate protein *in transit*.

The fact that the distribution of Fis1 in *mpf1Δ* cells could be completely corrected by higher levels of either Tom70 or Tom71 shows that the role of Mpf1 in regulating Fis1 is dispensable upon overexpression of one of these paralogues. This complementation can also suggest that Mpf1, Tom70, and Tom71 potentially share the same pathway in regulating the trafficking of Fis1 to mitochondria and peroxisomes. Further experiments to test whether Mpf1 directly interacts with Tom70 and/or Tom71 can shed light on the question if these proteins work together.

Although the precise molecular function of Mpf1 is not known yet, a hint for its physiological role is provided by our finding that yeast cells grown on fatty acid (oleate) benefit from the deletion of *MPF1*. We assume that this phenotype is due to an increase in the number of Fis1 molecules targeted towards peroxisomes, which subsequently enhance the number of peroxisomes, which could be particularly beneficial when oleate is the sole carbon source. However, we did not observe this effect on oleate upon deleting *TOM71 or TOM70* (data not shown), suggesting that other functions of Tom70/71 are still important when oleate is the sole carbon source. An alternative explanation is that in the absence of Mpf1 other cellular mechanisms (beside increased peroxisomal Fis1 levels) cause beneficial effects for growth in oleate media.

Another interesting aspect about Mpf1 is its rather short life-span. This inherent instability raises the question why cells produce protein molecules that will be degraded within minutes. Currently, we can only speculate that under some special conditions the presence of Mpf1 could be required immidiately and obtaining new molecules via enhancing transcription and translation might be too time consuming. In the future, it will be of interest to identify conditions that support enhanced stability of Mpf1.

Characterizing the sub-cellular localization of Mpf1 indicated it to be a peripheral membrane protein, loosely associated to the mitochondrial OM. Interestingly, even in the absence of both Tom70 and Tom71, Mpf1 still makes its way to the mitochondrial surface, suggesting the involvement of other factors that can mediate its association to the organelle. Mutating the conserved basic residues in the β1/β2 loop and thereby potentially disrupting the PH domain of Mpf1 did not hamper its mitochondrial localization either. Thus, it seems that either the triple mutation did not interfere completely with the function of the PH domain or other regions of the protein facilitate the association with mitochondria. Our finding that Mpf1(PH*) exhibited enhanced stability might indicate that certain proteins in the Ubiquitin/proteasome degradation pathway usually recognize a degron element in the PH domain but are unable to do so with the mutated PH domain. Alternatively, it might be that the mutations stabilize the interaction of Mpf1 with a protein and/or a lipid in the mitochondria outer membrane and through these interactions Mpf1 is stabilized.

Altogether, our findings contribute to novel insights on factors responsible for regulating the dual distribution of Fis1 to mitochondria and peroxisomes. We identified for the first time three proteins Tom70, its paralogue Tom71, and Mpf1 as involved in this process. In addition to recognizing a unique function of Tom71, we could provide a function for a so far uncharacterized protein – Mpf1. We identify the latter as an unstable protein at the surface of mitochondria. Collectvely, the current study provides the first glimpse into the process of dual distribution of TA proteins between mitochondria and peroxisomes.

## Materials and Methods

### Yeast strains and growth conditions

*Saccharomyces cerevisiae* strains used in this study are listed in Table S2. For induction of peroxisomes, yeast strains were grown at 30 °C on oleate-containing YNBO media (0.1% (w/v) yeast extract, 0.17% (w/v) yeast nitrogen base, 0.5% (w/v) ammonium sulphate, 0.0002% (w/v) uracil, 0.0002% (w/v) adenine sulphate, 0.12% oleic acid, 0.2% Tween40, supplied with amino acids). Generally, cells were grown at 30 °C on selective or rich media (YP) supplemented with 2% of either glucose or galactose. Yeast transformation was performed by the lithium acetate method (Gietz & Woods, 2006).

### Yeast growth assay

Yeast strains were cultivated till mid-logarithmic phase and after harvesting them, cells were resuspended to 1 ml of OD_600_ = 2. The cell suspension was fivefold serially diluted and 5 µl of each dilution was spotted on the indicated solid media. The plates were incubated at 30° C in a humid box and the growth was monitored for 2-10 days.

### High-throughput microscopy screening

The following query strains were made to cross with the yeast deletion library: (i) mCherry-Fis1, Pex3-GFP, (ii) mCherry-Fis1, Om45-GFP, (iii) mCherry-Gem1, Pex3-GFP, and (iv) mCherry-Gem1, Om45-GFP. To generate these query strains a DNA sequence encoding the mCherry tag was genomically inserted by homologous recombination at the 5’ of the sequence encoding either Fis1 or Gem1, with the strong and constitutive *TEF2* promoter and the Nourseothricin N-acetyl transferase (NAT) selection cassette. Subsequently, in these strains the DNA sequence encoding GFP was integrated by homologous recombination into the 3’ region of either Pex3 or Om45 also with the *TEF2* promoter and the hygromycin B phosphotransferase (HPH) selection cassette. Query strains were crossed by synthetic genetic array (SGA) with two libraries – the knock out (KO) library and the decreased abundance by mRNA perturbation (DAmP) library, as previously described (Cohen & Schuldiner, 2011; Tong & Boone, 2006). The high-throughput screen was performed by growing cells overnight at 30 °C in rich media (YP) supplemented with galactose, diluting them 1:10 in the next morning and letting them divide at 30 °C for 4 hours before imaging with an automated system (Breker et al., 2013).

### Recombinant DNA techniques

Full lists of primers and plasmids used in this study are found in Tables S3 and S4, respectively. The plasmid pGEM4-Mpf1-3HA was used as a template for site directed mutagenesis to create the PH domain mutant of Mpf1. The PCR product was digested with Dpn1 and transformed into *E. coli* cells. For gene deletion and manipulation, PCR product containing the selection cassette with flanking regions complementary to DNA sequences of the gene of interest were transformed into yeast cells by the Li-acetate method. Colonies were analysed by screening PCR. All constructs were verified by DNA sequencing.

### Separation of mitochondria and peroxisomes by gradient centrifugation

Yeast cells were precultured overnight in 100 ml YP medium supplemented with 0.1% glucose. Next morning, the culture was upscaled to 400 ml and incubated overnight. For induction of peroxisomes, cells were harvested (5000 x g, 6 min, RT) and washed with 20 ml sterile water and centrifuged again (5000 x g, 6 min, RT). Cells were then resuspended in 1000 ml YNBO medium and incubated for 16 – 20 hours. Then, cells were harvested (5000 x g, 6 min, RT) and washed twice with 30 ml sterile water followed by centrifugation (5000 x g, 6 min, RT). The cells were then incubated in 20 ml Dithiothreitol (DTT) buffer (100 mM Tris, 10 mM DTT) for 30 min and were harvested (1500 × g, 6 min, RT) and washed twice with 20 ml sorbitol buffer (20 mM 4-(2-hydroxyethyl)-1-piperazineethanesulfonic acid (HEPES), 1.2 M sorbitol, pH 7.2). Next, cells were incubated for 1 hour in 20 ml sorbitol buffer containing Zymolyase. Digestion of yeast cell walls was monitored by measuring the OD_600_ of small sample of cells to detect their rupturing upon addition of water.

All further steps were carried out on ice. Spheroplasts were washed twice with 20 ml sorbitol buffer and centrifuged (1500 × g, 6 min, 4 °C). Then, cells were homogenized using a dounce homogenizer in a solution of 15 ml lysis buffer (5 mM 2-(N-Morpholino)-ethane sulphonic acid (MES), 0.5 mM EDTA, 1 mM KCl) containing 0.6 M Sorbitol, Proteases inhibitors cocktail (PIC), 2 mM phenylmethylsulfonyl fluoride (PMSF), pH 5.5). Cell debris were removed by two centrifugation runs (1600 × g, 10 min, 4 °C). The resulting supernatant (containing mitochondria and peroxisomes) was centrifuged (13,000 × g, 5 min, 4 °C) and the pellet was resuspended in lysis buffer to OD_600_= 4. The organelles were loaded on top a density gradient consisting of 415 µl of 20%, 830 µl of 25%, 415 µl of 30%, and 830 µl of 40% Histodenz in gradient buffer A (5 mM MES, 1 mM EDTA, 1 mM KCl, and 0.1% (v/v) ethanol, pH 5.5). Gradients were centrifuged in a Beckman ultracentrifuge optima XE with a swinging bucket rotor, SW 60 Ti (100,000 × g, 90 min, 4 °C, acceleration 7, brake off). A total of 12 fractions with 235 µl in each were collected from the top of the gradient and mixed with 8x sample buffer (0.5 M Tris pH 6.8, 16% SDS, 80% glycerol, 8 mg/mL bromophenol blue) to a final 2x concentration. Then 5% (v/v) β-mercaptoethanol was added and the samples were heated at 95 °C. Fractions were subjected to SDS-PAGE followed by Western blotting.

### Isolation of Mitochondria

Yeast cells were grown in liquid media (volume of 2-6 L) to logarithmic phase. The cells were harvested (3000 x g, 5 min, RT), resuspended in DTT buffer and incubated at 30 °C for 15 min. Cells were harvested (2000 x g, 5 min, RT), washed once with spheroplasting buffer (1.2 M Sorbitol, 20 mM KPI, pH 7.2), harvested again and resuspended in spheroplasting buffer with Zymolyase (6 mg/g of cells) and incubated at 30 °C for 1 hour.

Further steps were carried out on ice. Spheroplasts were homogenized in homogenization buffer (0.6 M Sorbitol, 10 mm Tris, pH 7.4, 1 mM EDTA, 0.2% fatty acid-free BSA with 2 mM PMSF) using a dounce homogenizer to obtain a cell lysate. Cell debris and nuclei were removed by two clarifying spins (2000 x g, 10 min, 4 °C). The supernatant (cytosol + organelles) was centrifuged (18,000 x g, 15 min, 4 °C) to pellet crude mitochondria. The resulting post nuclear supernatant (PNS) consisted of ER/microsomal and cytosolic fractions. The crude mitochondria were washed twice with SEM buffer (250 mM Sucrose, 1 mM EDTA, 10 mM MOPS) containing 2 mM PMSF and were pelleted again (18,000 x g, 15 min, 4 °C).

### Subcellular fractionation

All the steps were carried out at 4 °C. Whole cell lysate and crude mitochondria were obtained as described above. To further purify mitochondria from potential contaminants, the mitochondrial fraction was layered on a Percoll gradient (25% Percoll, 2 M sucrose, 100 mM MOPS/KOH pH 7.2, 100 mM EDTA, 200 mM PMSF) and centrifuged (80,000 x g, 45 min, 4 °C, slow acceleration, slow brake). Highly pure mitochondria were found as a brownish layer close to the bottom of the tube and was removed carefully with a Pasteur pipette. The mitochondria were washed several times with SEM buffer containing 2 mM PMSF and pelleted again (18, 000 x g, 15 min, 4 °C).

To isolate ER/microsomal and cytosolic fractions, 20 ml of PNS was clarified (18,000 x g, 15 min, 4 °C) and centrifuged (200,000 x g, 1 hour, 4 °C). The supernatant contained the cytosolic fraction, and the brownish sticky pellet (consisting of ER) was resuspended in 2 ml of SEM buffer containing 2 mM PMSF and homogenized with a dounce homogenizer. The sample was centrifuged (18,000 x g, 20 min, 4 °C) to obtain ER/microsomes in the supernatant.

The obtained fractions were precipitated with chloroform-methanol mixture and the pellet was resuspended in 2x sample buffer (125 mM Tris pH 6.8, 4% SDS, 20% glycerol, 10% β-ME, 2 mg/mL bromophenol blue) to obtain protein concentration of 2 mg/ml. Samples were heated at 95 °C for 10 min and further analysed by SDS-PAGE and immunoblotting. Table S5 indicates the antibodies used in the current study.

### Proteinase K (PK) assay

Isolated mitochondria (100 µg) were incubated on ice for 15 minutes in 50 µL of either SEM buffer (untreated) or SEM buffer containing 10 µg/mL proteinase K (PK). Then, PK activity was inhibited by addition of 2 mM PMSF. The samples were centrifuged (18,000 x g, 15 min, 4 °C) and the pellets were resuspended in 2x sample buffer. Samples were heated at 95 °C for 10 min and further analysed by SDS-PAGE and immunoblotting.

### Carbonate (Alkaline) Extraction

Isolated mitochondria (100 µg) were resuspended on ice in 100 µL solution containing 20 mM HEPES, 2 mM PMSF and 1x PIC, pH 7.5. This was followed by the addition of 100 µL of carbonate solution 200 mM Na_2_CO_3_, 5 mM PMSF, 1x PIC) at various pH values (10.5, 11, 11.5, or 12), and further incubation for 20 min at 4 °C. Next, pellet (membrane proteins) and supernatant fraction (soluble proteins) were separated by centrifugation (75,000 x g, 30 min, 4 °C). The supernatant was precipitated by trichloroacetic acid (TCA). The pellet and precipitated proteins from the supernatant were resuspended in 40 µL 2x sample buffer, heated at 95 °C for 10 min and further analysed by SDS-PAGE and immunoblotting.

### Protein stability assay

Yeast strains were grown to mid-logarithmic phase. For each time point, cells corresponding to OD_600_ of 2 were collected and resuspended in 1 ml of media. Cycloheximide (CHX) at final conc. of 0.1 mg/ml was added at time=0 and the cells were incubated further at 30 °C for different time periods. Then, cells were harvested (3000 x g, 5 min, room temperature (RT)) and the proteins were extracted by alkaline lysis using 0.2 M NaOH, followed by heating with 2x sample buffer at 95 °C for 10 min. The samples were analysed by SDS-PAGE and immunoblotting.

### (Immuno) Fluorescence microscopy

Yeast cells were grown on synthetic media containing 2% glucose till mid-logarithmic phase. The cells (1 ml) were centrifuged (3000 x g, 5 min, RT) and the cells pellet was resuspended in 50 µl water. A small portion (5 µl) of this solution was mixed with 1% (w/v) low melting point agarose and was spread on a glass slide. Confocal spinning disc microscope was used to capture images and they were analysed using ImageJ (more details are given in the next section).

For immunofluorescene microscopy, a published protocol was optimized (Pemberton, 2014). Yeast cells were grown till mid-logarithmic phase and to fix them, they were incubated at 30 °C for 10 min with 1% (v/v) of 37% formaldehyde. Cells were washed twice and centrifuged (3000 x g, 5 min, RT) with Phosphate buffer (100 mM KH_2_PO_4_, 37.4 mM KOH, pH 6.5). Then, cells were resuspended in DTT buffer (100 mM Tris-HCl and 100 mM DTT) and incubated for 10 min at 30 °C. Cells were then washed twice with SPC buffer (1.2 M Sorbitol, 127 mM KH_2_PO_4_, 36 mM Citric acid) and spheroplasts were produced by incubating the cells for 45 min at 30 °C in SPC buffer + Zymolyase (6 mg/gram of cells). Spheroplasts were washed with SPC buffer and centrifuged (2000 x g, 5 min, 4 °C) and the pellets were resuspended in 100 µl SPC buffer, snap frozen, and stored at −80 °C.

Glass slides with 15 wells were treated with 0.1% (w/v) Poly-L-Lysine for 15 min at RT to enhance cell attachment. The poly-l-lysine was washed off by gently passing a stream of distilled water and the slides were air-dried. Next, 5 µl of spheroplasts solution were added to each well and were allowed to attach for 15 min. Excess liquid was removed, the slides were immersed in ice-cold MeOH for 5 min and were moved up and down 2-3 times. To further permeabilize the cell membrane, the slides were then immersed in acetone for 30 sec. Following this, the slides were air-dried and placed in a humid box for further steps. The cells were blocked with blocking buffer (PBS, 2% (w/v) milk, 0.1% (v/v) Tween-20) for 10 min at RT. The blocking solution was discarded, and cells were incubated at dark with 5 µg/ml primary antibody in the blocking buffer for 2 h at RT. Excess liquid was aspirated, and the cells were washed 3 times with PBS before mounting with 80% (v/v) glycerol. Cells were imaged using spinning disk microscope Zeiss Axio Examiner Z1 with a CSU-X1 real-time confocal system (Visitron) and SPOT Flex charge-coupled device camera (Visitron). Samples were observed using Zeiss Objective Plan-Apochromat 63×/1.4 Oil DIC M27. Images in Brightfield, GFP, and RFP channels were acquired through AxioVision software., Subsequent cropping and merging was done using Fiji software.

### Rapid protein extraction

Cells grown to mid logarithmic phase were harvested and resuspended such that 1 ml consisted of 2.5 units of OD_600_. The corresponding cell pellets were resuspended in 200 µl NaOH (100 mM) and incubated for 5 min at RT. The cells were centrifuged (3000 x g, 5 min, RT), resuspended in 2x sample buffer and heated for 5 min at 95 °C. The samples were centrifuged (3000 x g, 5 min, RT) and the supernatant was analysed using SDS-PAGE followed by immunoblotting.

### Pull-down assays

Cell pellets from 500 ml cultures were resuspended in 5 ml DTT buffer and incubated for 15 min in a 30 °C shaker. Cells were harvested (2000 x g, 5 min, RT), washed once with spheroplasting buffer (1.2 M Sorbitol, 20 mM KPI, pH 7.2), resuspended in spheroplasting buffer with Zymolyase (6 mg/g of cells), and then incubated for 1 hour in a 30 °C shaker. Cell pellets were resuspended in 1 ml lysis buffer (Tris-buffered saline (TBS), 2 mM PMSF, 1x EDTA free protease inhibitor cocktail, 5 mM EDTA) and homogenized using a douncer. Whole cell lysate (WCL) corresponding to 3 mg proteins was solubilized with 1% Triton X-100 and incubated in overhead shaker for 30 min at 4 °C. The solubilized sample was centrifuged (30000 g, 30 min, 4 °C) to clear out cell debris and the supernatant was incubated with anti-HA magnetic beads for 2 hours at 4 °C. The beads were washed with wash buffer (TBS, 0.5% Triton X-100, 5 mM EDTA, 350 mM NaCl) and the bound proteins were eluted by incubating the beads for 10 min at 55 °C with 2x sample buffer supplemented with 0.05% H_2_O_2_. Eluted material was supplemented with 5% β-ME, incubated at 95 °C for 5 min and analysed by SDS-PAGE followed by immunoblotting.

### Real-time quantitative PCR

RNA from 10 mL yeast culture was isolated using a mini kit for RNA isolation (Macherey-NAGEL, REF 740933.50). Next, 2.5 mg of RNA was used to prepare cDNA using the reagents and program mentioned in GoScript™ Reverse Transcription Mix, Oligo(dT) Protocol (Promega, A2791). RT-qPCR was set up in Thermo Fisher Scientific QuantStudio™ 5 Real-Time PCR System and the results were analyzed using Design and Analysis software 2.6.0. Primers used for the qPCR are listed in Table S3 and Actin was used as a reference.

## Acknowledgments

We thank E. Kracker for excellent technical assistance, H. Meyer and A. Fadel for help with the high-throughput screen, Marie Helmke and Luca Brenner for help with cloning, W. Girzalsky and R. Erdmann for anti-Pex14 antibody, and K.S. Dimmer for helpful discussions. This work was supported by the Deutsche Forschungsgemeinschaft (RA 1028/11-1 to D.R. and M.S.), the German-Israel Foundation (grant I-1458-412.13/2018 to M.S. and D.R.), and the Elisabeth and Franz Knoop-Foundation (fellowship to D.G.V.). All targeting work in the Schuldiner lab is also supported by an ERC CoG from the European Union (OnTarget, 864068). The robotic system of the Schuldiner lab was purchased through the kind support of the Blythe Brenden-Mann Foundation. MS is an Incumbent of the Dr. Gilbert Omenn and Martha Darling Professorial Chair in Molecular Genetics.

The authors declare no competing financial interests.

## Author contributions

N.A., D.G.V., designed and conducted experiments, J.O. performed experiments; E.Z., M.S., and D.R. designed experiments and analyzed data, N.A. and D. R. wrote the initial version of the manuscript. All authors read and contributed to the final manuscript.

